# Inflammatory and JAK-STAT Pathways as Shared Molecular Targets for ANCA-Associated Vasculitis and Nephrotic Syndrome

**DOI:** 10.1101/427898

**Authors:** Sean Eddy, Viji Nair, Laura H. Mariani, Felix H. Eichinger, John Hartman, Huateng Huang, Hemang Parikh, Jaclyn N. Taroni, Maja T. Lindenmeyer, Wenjun Ju, Casey S. Greene, Peter C. Grayson, Brad Godfrey, Clemens D. Cohen, Matt G. Sampson, Richard A. Lafayette, Nephrotic Syndrome Study Network (NEPTUNE), European Renal cDNA Bank – Else Kröner-Fresenius Biopsy Bank (ERCB), Vasculitis Clinical Research Consortium (VCRC), Jeffrey Krischer, Peter A. Merkel, Matthias Kretzler

## Abstract

**Background:** Glomerular diseases of the kidney are presently differentiated, diagnosed and treated according to conventional clinical or structural features. While etiologically diverse, these diseases share common clinical features including but not limited to reduced glomerular filtration rate, increased serum creatinine and proteinuria suggesting shared pathogenic mechanisms across diseases. Renal biopsies from patients with nephrotic syndrome (NS) or ANCA-associated vasculitis (AAV) were evaluated for molecular signals cutting across conventional disease categories as candidates for therapeutic targets.

**Methods:** Renal biopsies were obtained from patients with NS (minimal change disease, focal segmental glomerulosclerosis, or membranous nephropathy) (n=187) or AAV (granulomatosis with polyangiitis or microscopic polyangiitis) (n=80) from the <u>Nep</u>hrotic Syndrome S<u>tu</u>dy <u>Net</u>work (NEPTUNE) and the European Renal cDNA Bank. Transcriptional profiles were assessed for shared disease mechanisms.

**Results:** In the discovery cohort, 10–25% transcripts were differentially regulated versus healthy controls in both NS and AAV, >500 transcripts were shared across diseases. The majority of shared transcripts (60–77%) were validated in independent samples. Therapeutically targetable networks were identified, including inflammatory JAK-STAT signaling. *STAT1* eQTLs were identified and *STAT1* expression associated with GFR-based outcome. A transcriptional STAT1 activity score was generated from STAT1-regulated target genes which correlated with *CXCL10* (p<0.001), a JAK-STAT biomarker, predictors of CKD progression, interstitial fibrosis (r=0.41, p<0.001), and urinary EGF (r=-0.51, p<0.001).

**Conclusion:** AAV and NS caused from histopathologically distinct disease categories share common intra-renal molecular pathways cutting across conventional disease classifications. This approach provides a starting point for de novo drug development, and repurposing efforts in rare kidney diseases.

## Introduction

Rare diseases represent a unique challenge in both patient management and drug development. Nearly 30 million Americans, or approximately 10% of the population, are afflicted by at least one of over 7,000 rare diseases^1,2^. Because the patient population for any single rare disease cause can be quite small, understanding the underlying mechanisms of each disease and thus, the opportunity to develop and apply effective treatments is limited. As a consequence, the majority of rare diseases have no approved therapies^1,3^. With the advance of molecular profiling of tissue samples obtained from organ systems damaged by the rare disease, a data-driven understanding of the molecular basis across these diseases is in reach. Understanding shared molecular pathways activated during tissue damage can help broadly define mechanisms in distinct diseases that may be leveraged for drug discovery and repurposing across rare diseases^4^. By selectively targeting specific mechanisms of rare diseases responsible for chronic progressive end-organ dysfunction and damage, it may be possible to preserve organ function even when the disease initiating process is not directly impacted.

Nephrotic syndrome (NS) refers to a group of symptoms resulting from etiologically distinct diseases including focal segmental glomerulosclerosis (FSGS), minimal change disease (MCD), and membranous nephropathy (MN). To address the opportunity for drug repurposing and target identification in rare renal diseases, histologically distinct rare diseases (primary nephrotic syndrome (NS) and ANCA-associated vasculitis (AAV, granulomatosis with polyangiitis (GPA) and microscopic polyangiitis (MPA)) that each can result in end-organ kidney damage, were evaluated to identify shared molecular targets. Based on previous work that identified shared pathways across many distinct histopathologically defined chronic kidney diseases including primary forms of NS: FSGS, MCD and MN, as well as diabetic kidney disease and lupus nephritis^5^, the hypothesis tested was that transcriptional profiling in etiologically distinct rare diseases would enable the identification of shared disease mechanisms. Using primary NS (FSGS, MCD, and MN) and AAV as exemplar disease indications, we sought to test this hypothesis. These profiles were subsequently interrogated to identify shared activated pathways, with a focus on inflammation and fibrosis, and subsequently linked to clinical features and non-invasive biomarkers of pathway activation (Figure 1). Activated pathways were further evaluated to discover opportunities for drug repurposing, and identification of novel drug targets and/or biomarkers of kidney disease (Figure 1).

**Figure 1.**
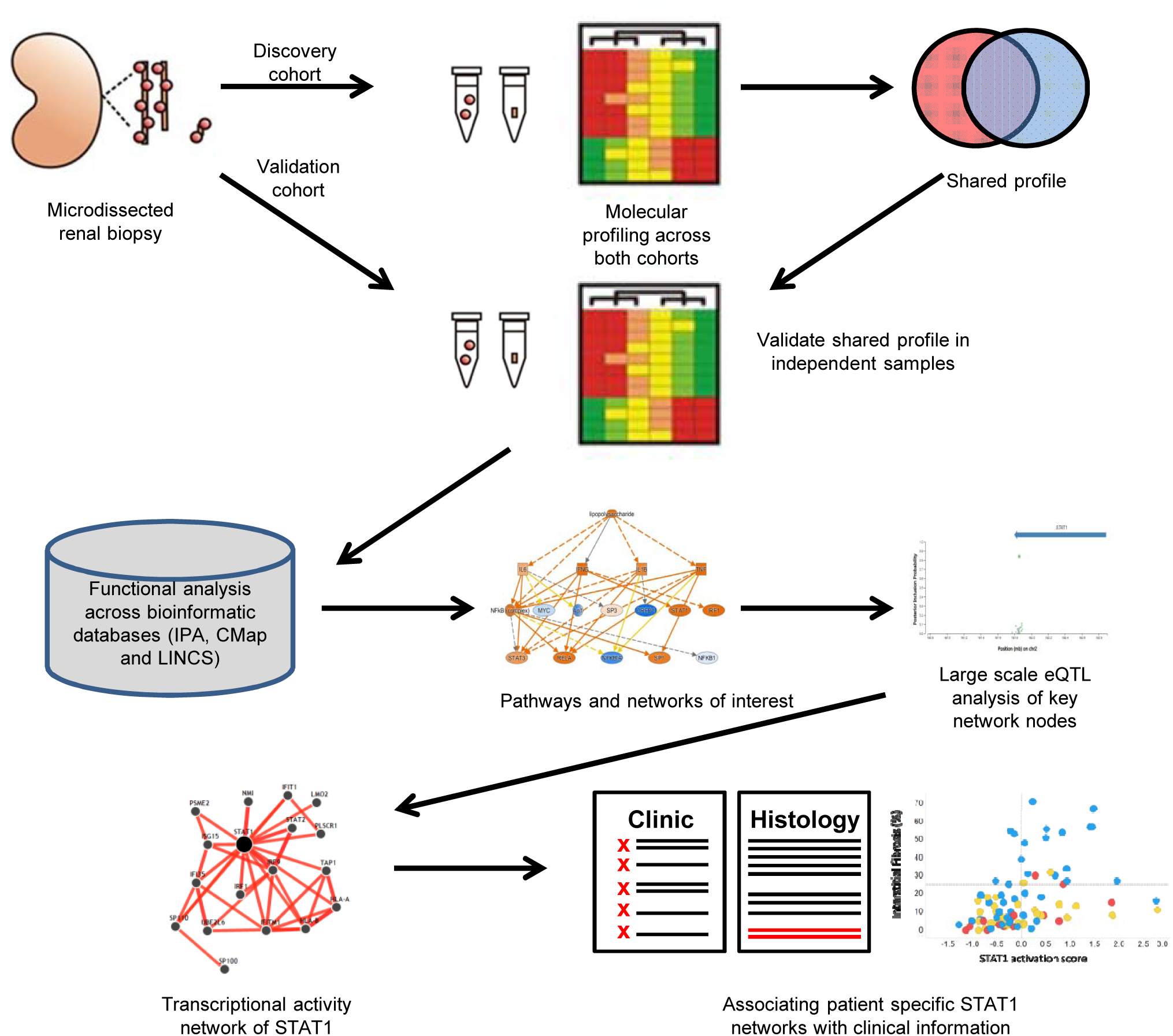
Schematic overview of the analysis approach for the approach identifying inflammatory targets and STAT1 activation in samples from subjects with nephrotic syndrome (NS) and ANCA-associated vasculitis (AAV).

## Results

Previous studies have indicated shared networks in FSGS, MCD, and MN^5^; and current studies from our group have demonstrated a continuum of expression between FSGS and MCD (unpublished data), we first sought to confirm common transcriptional programs in these primary forms of NS using data reduction and data-driven approach to assess transcriptional profiles from subjects with FSGS, MCD, or MN. Using transcriptional data generated from subjects with FSGS, MCD, and MN from ERCB, there were no distinct disease-specific clusters when a principal component analysis was performed on both the glomeruli or tubulointerstitium compartments (using data deposited in GEO under GSE104948 and GSE104954, Supplemental Figure 1a and 1b). To assess this statistically, eigen vectors for the first three principal components were assessed for differences across histopathologically classified diseases and displayed no evidence of significant differeces (Supplemental Figure 1c-e), thus FSGS, MCD, and MN were considered as a group of primary NS for comparison with AAV. To identify shared molecular mechanisms in NS and AAV (Figure 1), intrarenal transcripts in NS and AAV were evaluated in a discovery set from the ERCB. Affymetrix GeneChip–based steady state transcript expression from the kidney was separately derived from the glomeruli and tubulointerstitial compartment of kidney biopsies from 62 patients with NS and 23 patients with AAV from the ERCB (Table 1, Discovery cohort) and compared to intrarenal transcript profiles generated from 22 healthy living transplant donor biopsies (LD). In the tubulointerstitium, 1768 of the total of 11,666 expressed transcripts assessed in samples from patients with AAV and 565 genes in samples from patients with NS were differentially expressed (|FC|>1.3, q-value<0.05) versus LD. Of the 565 differentially expressed transcripts in the tubulointerstitium from subjects with NS, 90% (511/565) were also differentially expressed in the tubulointerstitium from subjects with AAV (Figure 2a). Importantly, the direction of expression was conserved across all transcripts in the tubulointerstitium, with a heatmap of the differentially expressed genes in the discovery cohort shown in Figure 2b. The top 10 differentially expressed transcripts in the tubulointerstitium from the discovery cohort are shown in Table 2, while a full list of differentially expressed transcripts is provided in Supplemental Table 1. In the glomeruli, 2322 genes were differentially expressed in AAV relative to LD, and 1059 in NS versus LD biopsies. A majority of differentially expressed transcripts in the glomeruli from patients with NS, (88% or 930/1059) were also differentially expressed in the glomeruli from patients with AAV, and nearly all (99.8% or 928/930) of shared transcripts were regulated in the same direction across diseases (Supplemental Figure 2, differentially expressed transcripts in the glomerular compartment are provided in Supplemental Table 2). These results are consistent with a common shared intra-renal transcriptional response across kidney diseases, independent of histolopathological diagnosis.

**Figure 2.**
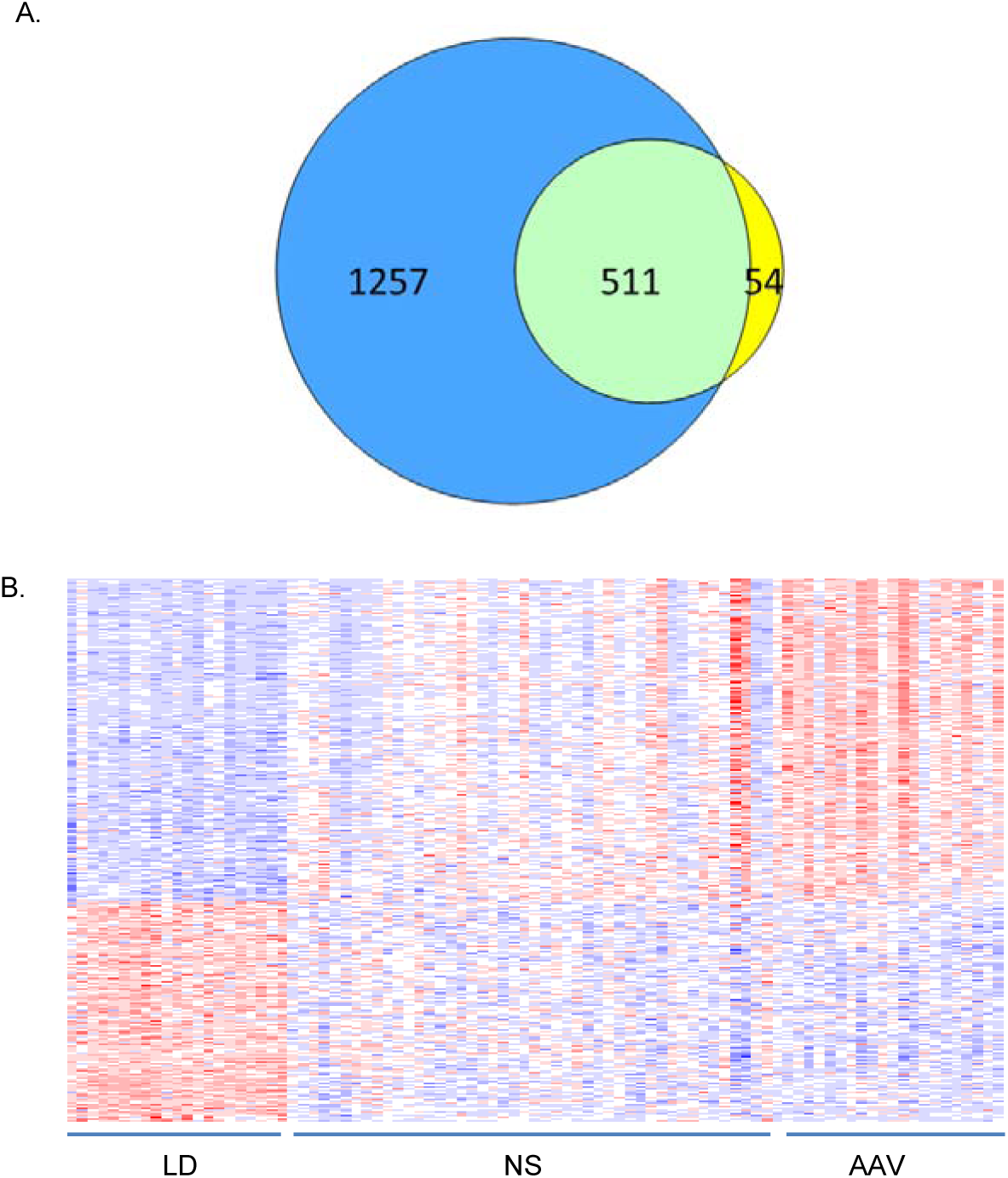
a.) Venn diagram of the number of tubulointerstitial differentially expressed genes (fold change>1.3 (up- and down-regulated genes), q-value<0.05) in ANCA-associated vasculitis (AAV) (blue circle) and nephrotic syndrome (NS) (yellow circle). b.) Heatmap representation of the 511 shared transcripts across samples in the dataset including living donors (LD). Expression values were plotted with row normalized Z-score scaling (blue = lower expression, red = higher expression).

**Table 1.** Demographic information of subjects profiled in the discovery and validation cohorts. Age and eGFR are reported as mean ± standard deviation.

| Discovery cohort | Nephrotic syndrome | ANCA-associated vasculitis | Living donor |
| --- | --- | --- | --- |
| Glom samples | 58 | 22 | 21 |
| TI samples | 47 | 21 | 21 |
| eGFR, MDRD<br>(ml/min/1.73m<sup>2</sup>) | 83 ± 39 | 46 ± 31 | 104 ± 31 |
| Age | 47.2 ± 17.7 | 58.0 ± 13.8 | 47.3 ± 11.5 |
| Sex<br>(%Female) | 45 | 43 | 45 |
| Validation Cohort |  |  |  |
| Glom samples | 90 | 15 | 6 |
| TI samples | 107 | 57 | 5 |
| eGFR, MDRD<br>(ml/min/1.73m<sup>2</sup>) | 77 ± 32 | 31 ± 27 | 108 ± 33 |
| Age | 47.1 ± 15.7 | 57.8 ± 9.8 | 52.5 ± 6.9 |
| Sex<br>(%Female) | 33 | 58 | 57 |

**Table 2.** Top 10 differentially expressed genes sorted by the average absolute fold change across both AAV and NS relative to living donors.

| Gene Symbol | Entrez GeneID | AAV vs. LD log2 fold-change | AAV vs. LD q-value | NS vs LD log2 fold-change | NS vs LD q-value |
| --- | --- | --- | --- | --- | --- |
| LTF | 4057 | 4.73 | <0.0001 | 3.59 | <0.0001 |
| CXCL6 | 6372 | 2.92 | <0.0001 | 2.81 | <0.0001 |
| LYZ | 4069 | 3.13 | <0.0001 | 2.15 | <0.0001 |
| REG1A | 5967 | 2.87 | <0.0001 | 2.25 | <0.0001 |
| OLFM4 | 10562 | 2.77 | <0.0001 | 1.71 | 5.4E-3 |
| NNMT | 4837 | 2.91 | <0.0001 | 1.38 | 2.2E-4 |
| SERPINA3 | 12 | 2.94 | <0.0001 | 1.33 | <0.0001 |
| C3 | 718 | 2.72 | <0.0001 | 1.48 | <0.0001 |
| PDK4 | 5166 | -1.97 | <0.0001 | -2.12 | <0.0001 |
| MMP7 | 4316 | 2.47 | <0.0001 | 1.56 | 3.4E-4 |

Hierarchical clustering was then performed on the differentially expressed genes in the discovery cohort to explore sample and cluster level similarities or differences in expression profiles from patients with nephrotic syndrome and ANCA-associated vasculitis. Using the tubulointerstitial data, hierarchical clustering identified four clusters (Figure 3a). The first cluster contained all of the healthy LD samples, as well as a broad spectrum of samples across kidney diseases (MCD, FSGS, AAV), while clusters 2, 3, and 4 contained samples from all diseases, but no LD (Figure 3b). The association of transcript clusters with phenotypes was evaluated by assessing eGFR across the clusters (Figure 3c). Within each histological diagnosis, significant eGFR differences were observed across clusters (ANOVA, p<0.05) (Figure 3c). The majority of transcriptional profiles from MCD patient samples were found in clusters 1 and 2 and these were from patients with MCD that had the highest mean eGFR (110±20 and 82±35 in clusters 1 and 2, respectively). Conversely, a majority of profiles from AAV patient samples were found in clusters 3 and 4 (Figure 3b), and were from patients with the lowest eGFR across AAV (14±10 and 15±10 for clusters 3 and 4 respectively). Samples from patients within each histological diagnosis, had significantly lower eGFR in cluster 4 (p<0.001), and increased transcripts in this cluster of patients were negatively correlated with log2 eGFR across diseases. For example, the top differentially regulated genes in cluster 4 - *CXCL6, C3* and *SERPINA3* showed a negative GFR correlation (r≤-0.4, p<0.001 for all three genes, Figure 3d) linking an inflammatory disease state to impaired renal function independent of underlying histological diagnosis.

**Figure 3.**
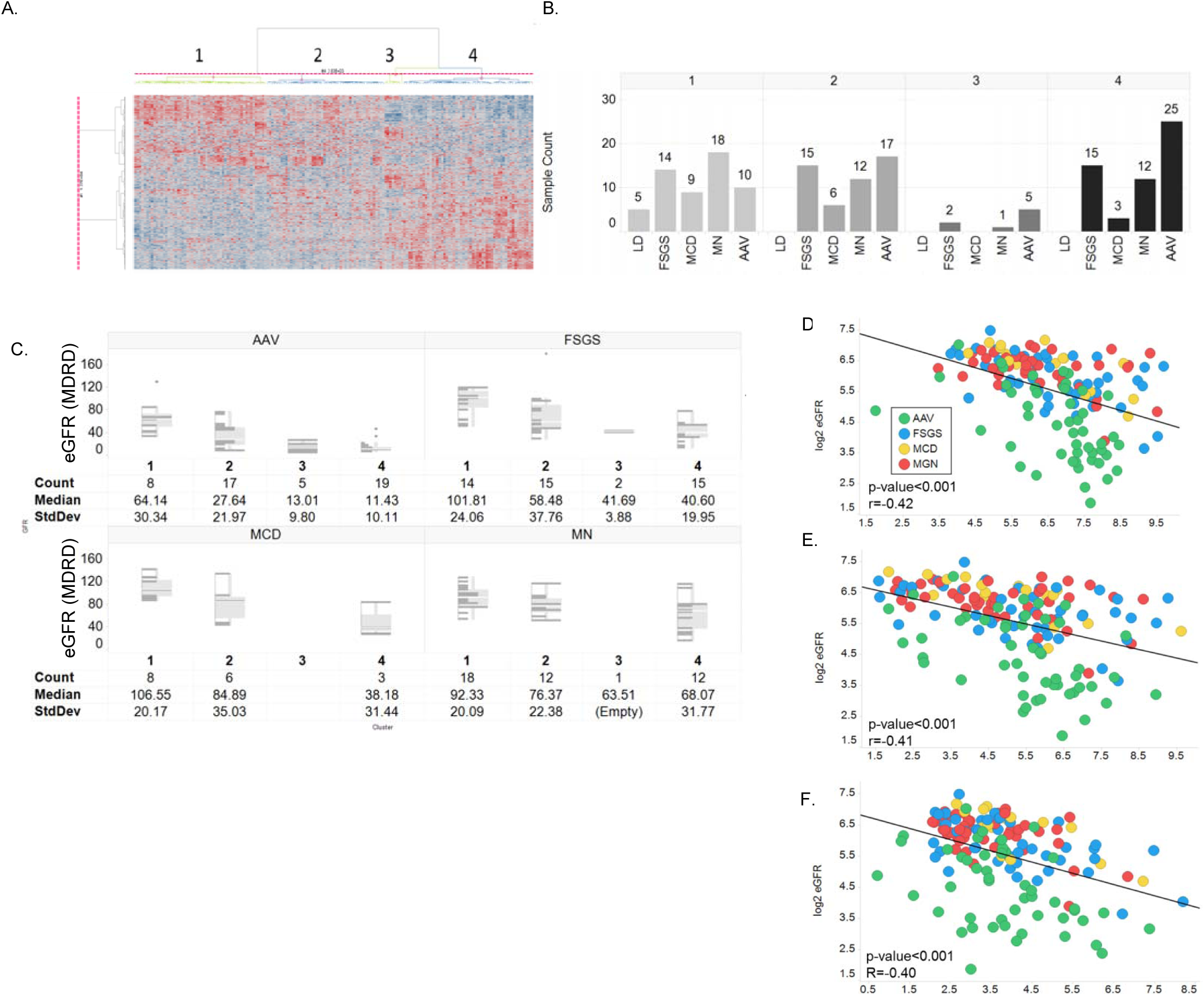
A.) Hierarchical clustering (Ward’s method) of shared transcripts identified in the tubulointerstitial discovery set on the test data set. B.) Bar chart of histological disease diagnosis, focal segmental glomerulosclerosis (FSGS), minimal change disease (MCD) and membranous glomerulonephropathy (MN), AAV in each of the four clusters. C.) Box plots with data distribution of eGFR (ml/min/1.72m^2^) by histological diagnosis and cluster. D.) Correlation of log2 mRNA expression with log2 eGFR of select transcripts enriched in patients within cluster 4: D. *C3*, E. *SERPINA3*, and F. *CXCL6* (right panel) enriched in patients in cluster 4 and correlated with eGFR across the entire cohort.

To validate shared transcripts in the tubulointerstitium and glomeruli, we profiled an independent set of samples for AAV from ERCB and an independent set of samples for nephrotic syndrome from NEPTUNE on Affymetrix ST2.1 gene chips (Table 1, Validation cohort), assessing the 511 shared transcripts identified in the tubulointerstitium and the 928 shared transcripts in the glomeruli from the discovery set (Table 3). In the tubulointerstitium, 508 of 511 genes identified in the discovery cohort were expressed above background expression levels in the validation cohort and could be assessed for differential expression, while all 928 differentially expressed genes in the glomeruli were available for assessment in the validation cohort. Table 3 provides summary level information and individual genes validated in both the tubulointerstitium and glomeruli; full results are provided in Supplemental Tables 3 and 4 for each compartment, respectively. Thus, the vast majority of transcripts identified as differentially regulated and shared between diseases in the discovery phase were validated in a test dataset of independent samples.

**Table 3.** Summary level information for shared gene expression profiles in the glomeruli and tubulointerstitium from the discovery and validation cohorts. In the discovery cohort, differentially expressed (DE) genes were defined as those with | FC | >1.3 and q-value<0.05.

| Discovery cohort | TI | Glom |
| --- | --- | --- |
| AAV DE genes | 1768 | 2322 |
| NS DE genes | 565 | 1059 |
| Common DE genes (AAV + NS) | 511 | 928 |
| **Validation cohort** |  |  |
| Common DE genes present in validation cohort | 508 | 928 |
| Common DE genes (AAV + NS) $q < 0.05$ | 308/508 (60%) | 592/928 (64%) |
| Common DE genes (AAV + NS) $q < 0.1$ | 396/508 (77%) | 609/928 (66%) |

The complement gene, *C3* was among the highest up-regulated transcript in both tubulointerstitial and glomerular samples from patients with NS and AAV in the discovery and validation cohorts (Figure 4a-d). As complement activation has been implicated in AAV^6^, the status of the pathway in AAV and NS was investigated further. Multiple genes in the complement pathway showed tissue level transcript up-regulation in both NS and AAV in the TI (Figure 4e). The complement system pathway was one of the most significantly increased canonical pathways as tested by a panel of enrichment strategies (Table 4).

**Figure 4.**
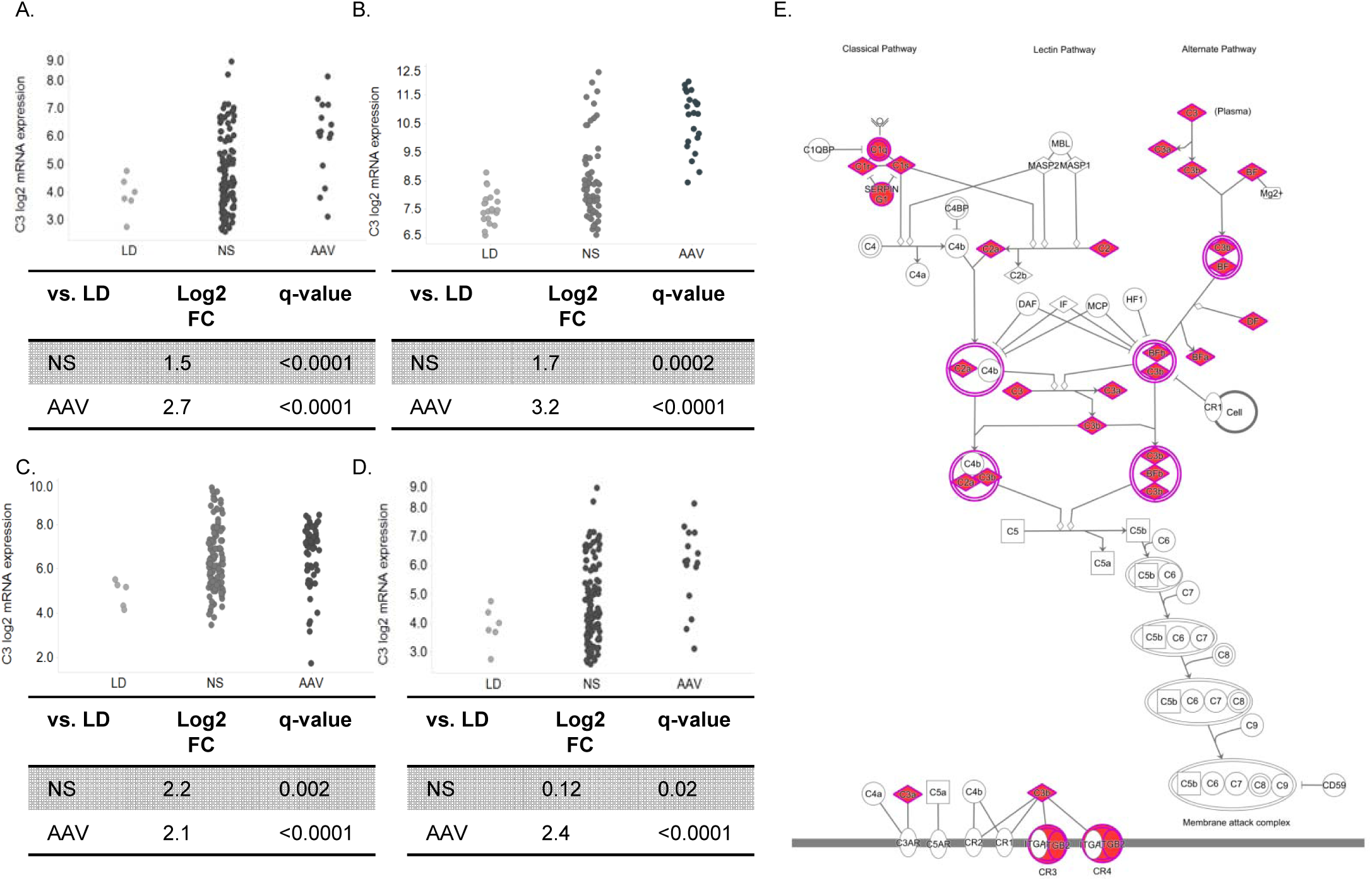
mRNA expression profile of *C3* from samples from LD, and NS and AAV in the: A.) tubulointerstitium from the discovery cohort, B.) glomeruli from the discovery cohort, C.) tubulointerstitium from the validation cohort, D.) glomeruli from the validation cohort. E.) Complement system pathway diagram indicating the genes in the complement pathway up-regulated (colored red) in the tubulointerstitium of diseased samples in both the discovery and validation cohort.

**Table 4.** Significant enrichment of complement pathways in IPA and Genomatix Genome Analyzer from the 396 and 609 shared and validated transcripts in the TI and glomeruli, respectively.

| Compartment | Canonical Pathway | Source | Enrichment p-value |
| --- | --- | --- | --- |
| TI | Complement System | IPA Canonical Pathways | 9.68E-7 |
| TI | Classical Complement Pathway | GGA | 3.48E-4 |
| TI | Alternative Complement Pathway | GGA | 1.73E-2 |
| Glomeruli | Complement System | IPA Canonical Pathways | 4.38E-6 |
| Glomeruli | Classical Complement Pathway | GGA | 6.50E-3 |
| Glomeruli | Alternative Complement Pathway | GGA | 2.65E-2 |

While individual gene expression profiles can be identified and mapped as drug targets, identifying the upstream mechanisms influencing downstream gene expression is likely a more rigorous strategy to identify functionally active, and potentially targetable signaling networks^7–9^. To derive the activation state of targetable pathways from the transcriptional profiles, two approaches were taken. The first strategy assessed whether any of the validated differentially expressed transcripts (396 shared transcripts with q-value<0.1 in the tubulointerstitium) could be explained by predicted upstream regulators. This was paired with an independent approach to match the shared transcriptional profile with those of cell lines exposed to various perturbagens targeting known pathways (small molecules, over-expressing constructs, and siRNA knockdown).

Upstream regulators are molecules (including but not limited to: transcription factors, growth factors, kinases, chemicals) predicted to affect activity of downstream expression profiles. Activities are predicted using literature derived cause and effect relationships, e.g. up-regulation of a transcription factor target gene supports a predicted activation in the transcription factor, while down-regulation supports inhibition^7–9^. The top biological networks from upstream regulator analysis in the TI are shown in Table 5 (an expanded list of results is provided in Supplemental Table 5; glomerular results are found in Supplemental Table 6). Predicted activities of upstream regulators are based on directionality of underlying gene expression that is similar to (predicted activation) or inverse (predicted inhibition) of the uploaded differential expression signature. The top enriched upstream regulators were indicative of inflammatory pathway activation, and included predicted activation of cytokines IFNγ, TNF and predicted inhibition of the glucocorticoid receptor (NR3C1). A mechanistic network of interconnected upstream regulators was discovered (upstream regulators that can explain multiple downstream transcripts as part of a network) (Figure 5a), and explains 56% of the downstream gene expression changes in the shared, tubulointerstitial data set ((222/396)).

**Figure 5.**
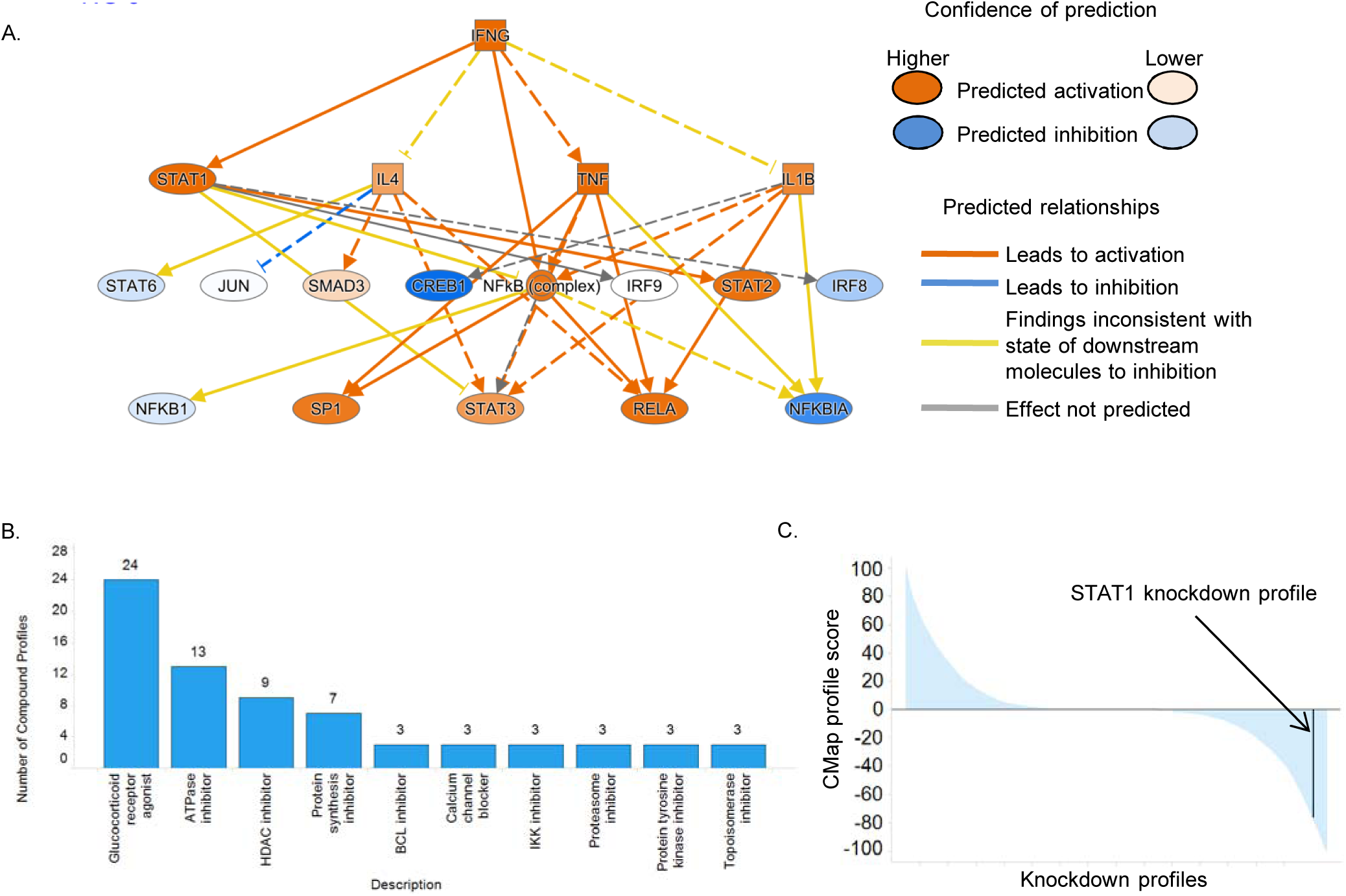
A.) Inflammatory mechanistic network from IPA upstream regulator analysis. B.) Bar chart indicating the number of compounds in the indicated compound classes with CMap profile scores ranging from -50 to -100. C.) Waterfall plot of CMap profile scores by knockdown experiments, with the STAT1 knockdown experiment indicated.

**Table 5.** Ingenuity Pathways Analysis Results for the top 15 biological network nodes (by p-value enrichment) with a prediction of activated or inhibited based on underlying gene expression profiles.

| Upstream Regulator | Molecule Type | Predicted Activation State | Activation z-score | p-value of overlap |
| --- | --- | --- | --- | --- |
| IFNG | cytokine | Activated | 4.32 | 4.23E-34 |
| TNF | cytokine | Activated | 2.94 | 2.42E-32 |
| NR3C1 | ligand-dependent nuclear receptor | Inhibited | -2.69 | 1.80E-16 |
| OSM | cytokine | Activated | 2.81 | 4.16E-16 |
| PDGF BB | complex | Inhibited | -3.56 | 1.08E-15 |
| Vegf | group | Inhibited | -2.29 | 5.66E-14 |
| CREB1 | transcription regulator | Inhibited | -2.26 | 2.97E-13 |
| MAPK1 | kinase | Inhibited | -3.39 | 1.15E-12 |
| IL1 | group | Activated | 2.11 | 1.52E-12 |
| IRF1 | transcription regulator | Activated | 3.39 | 3.86E-12 |
| Pkc(s) | group | Inhibited | -2.11 | 9.64E-12 |
| STAT1 | transcription regulator | Activated | 3.62 | 2.90E-11 |
| FSH | complex | Inhibited | -2.44 | 8.75E-11 |
| Interferon alpha | group | Activated | 4.21 | 9.89E-11 |
| IL27 | cytokine | Activated | 3.39 | 1.63E-10 |

To further define functional associations, the 396 gene shared transcriptional profile from the tubulointerstitium was used to query Connectivity Map (CMap) L1000 profiles^10^. The resulting CMap output contains a ranking of experimentally generated transcriptional profiles generated in cell lines that are inverse (potential therapeutic) or similar (potentially exacerbating disease) to the shared transcriptional profile. This provided an additional, independent, data-driven approach to identify signaling networks and potential drug targets active in patients with NS and AAV. For these analyses, CMap profiles were limited to those with absolute profile scores >50 (13% of CMap profiles met this criteria). Looking at compound profiles that are inversely related to the shared signature, profiles of glucocorticoid agonists were the most frequent (Figure 5b). Calcium channel blockers were also identified. IKK inhibitors, which block NFκB activation were also identified as inversely related to the shared signature. Activation of NFκB and RELA (p65 subunit of NFκB) were also predicted as part of the shared transcriptional profile and are present in the upstream regulator network (Figure 5a). Interestingly, a number of profiles from ATPase inhibitors, HDAC inhibitors, protein synthesis inhibitors, and proteasome inhibitors were also identified. Over-expression (OE) experiments (Supplemental Table 7) with the highest CMap profile scores indicating a signature similar to the shared molecular profile included HLA-DRA (99.91), consistent with an inflammatory signal and PSMB10 (95.43), which would suggest elevated proteasome signaling shared across diseases.

To gain further insight into key signaling networks shared across diseases, upstream regulators with predicted activities from IPA were compared with CMap profiles to test for consensus between these two independent modeling approaches. The top 15 unique TI networks are shown in Table 6, (multiple CMap profiles for a compound are often observed due to profiles being generated for different treatment concentrations or duration) with 67% (10/15) showing consensus using independent approaches. These included networks representing lower glucocorticoid signaling (dexamethasone and fluticasone), reduced transcription factor activities in the TI of FOXA1 and HNF4A, and activation of CD44, STAT1, and SOX2 (Table 6). In CMap profiles, experimental knock-down of STAT1 resulted in an expression profile that was the inverse of the shared disease profile, and was among the top 5% of profiles showing this inverse relationship (Figure 5c). In the glomeruli, 18 unique nodes were identified (Supplemental Table 8) and 61% (11/18) were consistent across the two analyses. These included nodes representing a lowered glucocorticoid signal (dexamethasone), reduced transcription factor activities of FOXA1, HNF1A and HNF4A, and activation of TLR7, IL2, CCND1, PLAUR, IFNGR1, and NR0B2. These pathways are consistent with inflammatory activation and decreased epithelial differentiation states common among NS and AAV.

**Table 6.**
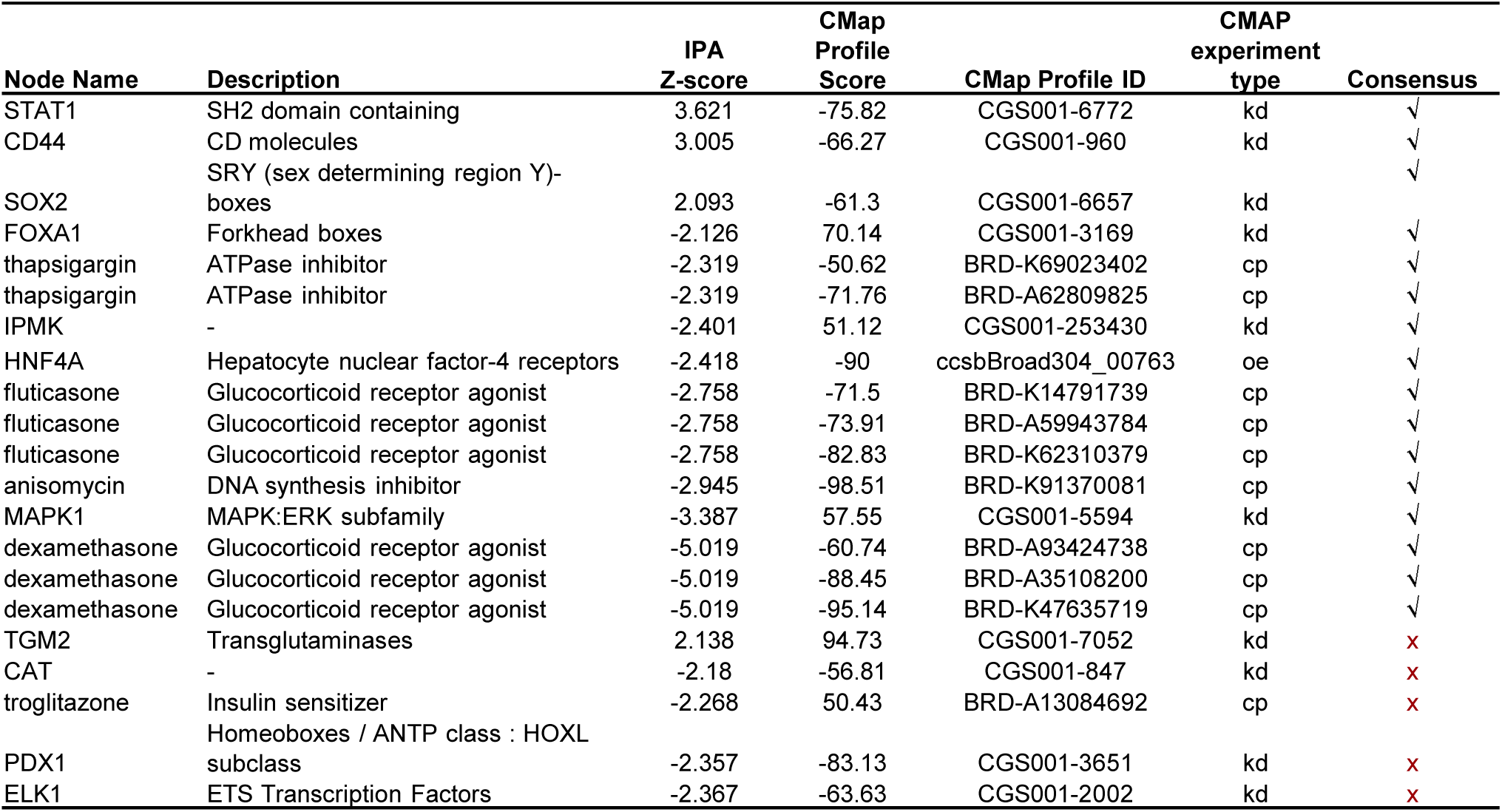
IPA nodes with corresponding profile scores from CMap and LINCS database (CLUE) using the shared TI transcriptional profile.

Consensus findings from IPA and CMap profiles were then investigated for the potential underlying genetic components that might contributed to an alteration in disease-associated expression and activity. TI and glomerular-specific cis-eQTLs were generated across the NEPTUNE cohort (www.nephqtl.org)^11^; cis-eQTLs were fine mapped using the Bayesian Deterministic Approximation of Posteriors^12^, which identifies clusters of variants predicted to drive eQTL associations. *STAT1*, *CD44*, *SOX2*, *FOXA1*, *ATP2A1*, *ATP2A2*, *ATP2A3* (the ATP2A family members encode sarcoplasmic reticulum Ca^2+^ ATPase and are targets of thapsigargin), *IPMK, HNF4A*, *MAPK1*, and *NR3C1* (translated protein is the glucocorticoid receptor, a target of prednisone and dexamethasone) were assessed in TI-specific eQTL analyses, and *TLR7*, *IL2, NR0B2*, *PLAUR*, *IFNGR1*, *FLT3* (the translated protein is a target of sunitinib) *FOXA1*, *HNF4A*, *HNF1A*, and *NR3C1* were assessed in glomerular-specific multi-SNP eQTL analyses. Analyses of eQTLs for these genes in the TI and glomeruli only identified STAT1 with a gene level FDR<0.05 (Supplemental Table 9. As variants associated with *STAT1* expression were non-coding, expression of *STAT1* was investigated for associations with outcome across the NEPTUNE cohort. Samples were binned in tertiles according to *STAT1* expression. In an unadjusted model, elevated *STAT1* expression was significantly associated with faster progression to a composite eGFR endpoint (time to ESRD or 40% loss of eGFR) (log rank p-value<0.01) (Figure 6).

**Figure 6.**
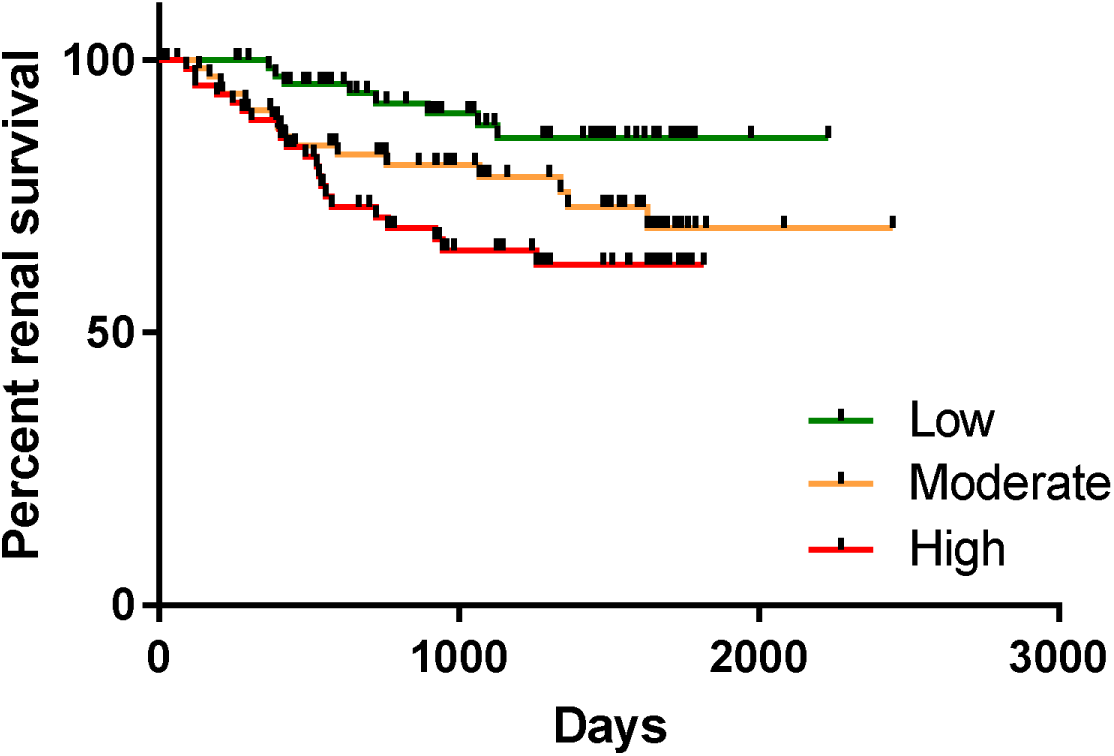
Kaplan-Meier analysis of tubulointerstitial *STAT1* expression and its association with outcome in the NEPTUNE cohort.

Because expression of a single gene is not necessarily indicative of network or pathway activity, a STAT1 score was generated, given that a similar approach yielded a JAK-STAT activation score that was associated with increased STAT1 and STAT3 phosphorylation in FSGS^13^. Using a data driven approach, a kidney-specific STAT1 network comprised of 17 genes was identified^14^, and further refined for genes functionally up-regulated by IFNγ stimulation by utilizing the Database of IFN Regulated Genes, Interferome^15^. Transcripts were considered validated if they showed at least 2-fold increase by IFNγ stimulation in at least 5 datasets. Next, the genes were assessed for the presence of STAT1 or STAT family binding sites, bringing the STAT1 activated gene set to 14 genes (detailed in Supplemental Table 10). The STAT1 activation score was strongly correlated with STAT1 expression in the discovery (r=0.95, p<0.001, n=68) and validation cohorts (r=0.89, p<0.001) (data not shown). IP-10 (CXCL10) has been used as a target engagement and renal inflammation biomarker in a clinical trial assessing the effectiveness of JAK inhibition in patients with diabetic kidney disease treated with baricitinib^16^, but was not used to generate the STAT1 activation score. The STAT1 activation score strongly correlated with *CXCL10* mRNA levels in both the discovery (Figure 7a, r=0.81, p<0.001, n=68) and validation (Figure 7b, r=0.58, p<0.001, n=166) AAV and NS datasets, as it did in DKD in confirmation of the clinical trial observation^17,18^ (Supplemental Figure 3). Because STAT1 activation can be considered a reflection of the intra-renal inflammatory state and has been reported to drive tissue fibrosis^19^, the interaction of the STAT1 activation score with interstitial fibrosis was assessed. In NEPTUNE NS patients, the STAT1 activation score was correlated with interstitial fibrosis (Figure 7c, r=0.41, p<0.001, n=95). Similar results were found associating STAT1 activation score with tubular atrophy (Supplemental Figure 4). Intrarenal epidermal growth factor *(EGF)* mRNA levels and urinary EGF to creatinine ratios (uEGF) have been identified as markers of tubular differentiation and regeneration. Indeed, renal tissue *EGF* mRNA and urinary EGF protein have been reported to negatively correlate with interstitial fibrosis and tubular atrophy, and can predict progression of CKD^17^. The STAT1 activation score significantly and inversely correlated with *EGF* mRNA in both the discovery and test datasets (Figure 7d, r=-0.58, p<0.001, n=68 and Figure 7e, r=-0.57, p<0.001, n=166), and uEGF (Figure 7f, r=-0.51, p<0.001, n=62). In each case the STAT1 activation score was negatively correlated with *EGF* and uEGF, supporting the link of inflammation, epithelial dysfunction, renal fibrosis and long term progression of glomerular disease.

**Figure 7.**
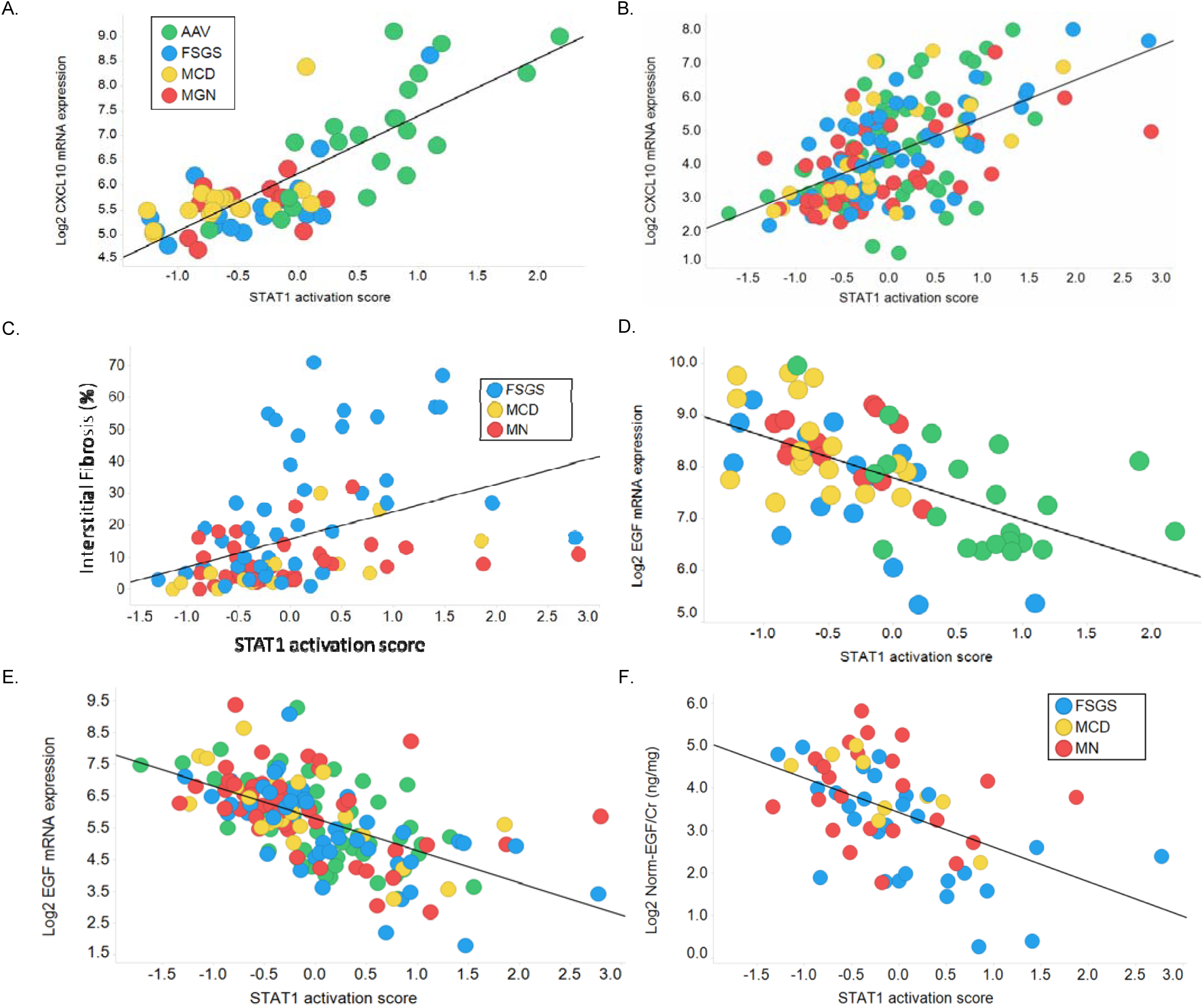
Association of STAT1 activation scores with biomarkers and clinical data. Correlation of the STAT1 activation score with *CXCL10* mRNA in the A.) Discovery cohort and B.) Validation cohort, with C.) interstitial fibrosis from the NEPTUNE cohort, with *EGF* mRNA in the D.) Discovery cohort and E.) Validation cohort, and with F.) urinary EGF from the NEPTUNE cohort.

## Discussion

Parenchymal organs require high degrees of differentiation to perform their specialized functions. This differentiation, however, can limit cellular programs that are activated in response to a diverse set of injury mechanisms. Tissues require, at least in part, dedifferentiation to carry out cellular damage responses^20–22^. Across many epithelial organs a uniform response consisting of loss of epithelial function, innate immune response activation, and activation of chronic fibrotic responses can be observed^23–26^. Through the evaluation of shared tissue injury mechanisms across diseases, pathways associated with progressive loss of organ function may be used to identify disease-modifying pathways and networks as therapeutic targets capable of impacting patient care in multiple rare diseases. In the setting of the NIH Rare Clinical Disease Research Network this hypothesis was tested by mapping the transcriptional profiles of different diseases that cause kidney damage.

Consistent with previous findings of shared kidney-specific transcriptional mechanisms among patients with chronic kidney disease from various causes, in particular a strong transcriptional overlap in found in patients with diabetic nephropathy, lupus nephritis, and IgA nephropathy^5^, an analysis of samples from patients with diverse causes of primary NS or AAV yielded a large number of shared transcripts and mechanisms that were shared across conventional histological disease classifications. The majority of transcripts differentially expressed in NS (90%) were also found to be differentially expressed in AAV, were directionally consistent, and were validated in an independent set of patient samples. Of note, multiple genes in the classical and alternative complement pathway were among the shared altered transcripts in AAV and NS, in both the TI and glomeruli. Given the role that the alternative complement pathway plays in the kidney of patients with AAV^6^, Phase II trials suggesting the efficacy of alternate complement activation in patients with AAV through inhibition of C5a^27^, and the heterogeneous *C3* mRNA expression upstream of the alternative pathway, expanded targeting of complement activation in patients with other kidney diseases may be warranted.

Using the validated gene set for further analyses, a convergence of molecular mechanisms was identified including activation of inflammatory pathways and suppression or inhibition of differentiation related pathways. For the inflammatory-related pathways, evidence for their potential causal role in autoimmune or renal disease has been established, including TNF^28–30^, JAK-STAT^16^, NFκB^31^, proteasome signaling^32,33^, and re-activation of glucocorticoid signaling through the use of glucocorticoid agonists^34^. JAK-STAT pathway activation has already been implicated across other kidney diseases including diabetic kidney disease (DKD)^35–38^, and FSGS^13^. In addition to the inflammatory-related pathways, reduced HNF4 transcriptional activity could represent a state of de-differentiation as Hnf4a can induce mesenchymal to epithelial transition^39^. Hnf4a co-expressed with Emx2, Hnf1b, and Pax8 has been shown to induce tubular epithelial cell differentiation^40^. Furthermore, HNF4A was identified as a hub protein in a protein-protein interaction network of CKDGen genes with eGFR-associated expression, suggesting that reactivation of the HNF4A transcriptional program could have broad implications across CKD^5^.

Expertly curated causal inference associations, pathway enrichment, and experimentally-derived computational approaches to identify perturbations associated with a shared transcriptional profile in NS and AAV all converge on transcriptional mechanisms related to inflammatory and immune signaling, JAK-STAT, complement activation, and HNF4A. An assessment of converagent mechanism in the compartment specific eQTL landscape in kidney disease was performed across the NEPTUNE cohort. This analysis identified clusters of non-coding snps that were associated with *STAT1* expression in the tubulointerstitium, and elevated *STAT1* expression was associated with faster progression to a composite eGFR endpoint (40% loss of eGFR or ESRD).

Because transcription factor activation is driven by its ability to activate downstream transcription and not simply expression, we generated a STAT1 activation score. This approach was previously used to characterize JAK-STAT pathway activation and correlated with activated (phos-STAT) activation in FSGS^13^ in the NEPTUNE cohort. The STAT1 score correlated across diseases on an individual patient level with *CXCL10* (IP-10) mRNA, and urinary EGF. IP-10 has been used as a pharmacodynamic biomarker in a Phase II trial suggesting the efficacy of baricitinib, an inhibitor of JAK1 and JAK2, in diabetic kidney disease^35^. Given the successful phase II trial of baricitinib in DKD, the similarity of the STAT1 score in AAV relative to DKD, and the superiority of baricitinib compared to TNF inhibition in a distinct systemic autoimmune disease, rheumatoid arthritis ^41^, argues for clinical evaluation of JAK inhibitors in patients with AAV. In addition, the similar level of signaling identified in NS, along with activation being associated with a progression biomarker, lends support to developing baricitinib for treatment of NS as well. Using a biomarker-driven approach in a precision medicine strategy, patients with active JAK-STAT signaling regardless of histological diagnosis could potentially benefit from targeted inhibition of the pathway using a basket clinical trial design^42^. The FDA recently approved the first treatment for cancer that applies a common biomarker to guide decisions about therapy, cutting across solid tumors as opposed to approval based on histological assessment of tumor origin.^43^

This work establishes the landscape of underlying molecular mechanisms shared between nephrotic syndrome and ANCA-associated vasculitis. In a more general sense, identifying common disease mechanisms shared across rare diseases is an approach that may expand the currently limited treatment options. This may help to overcome limits in the molecular understanding of rare diseases, identify novel and known targetable mechanisms shared across diseases, and use the knowledge towards therapeutic repurposing for available drugs, ultimately improving treatment options for patients.

## Methods

### Patient cohorts

For the discovery cohort, samples from adult patients with AAV (n=23) and NS (n=62) were used from the European Renal cDNA Bank (ERCB). ERCB is a European multicenter study that collected renal biopsy tissue for gene expression analysis in RNA fixative (RNAlater, Qiagen), coupled with cross-sectional clinical information collected at the time of a clinically indicated renal biopsy^44^. For the validation cohort, samples from adult patients with AAV (n=57) were obtained from the ERCB, and samples and clinical data from adult patients with NS (n=116) were obtained from the Nephrotic Syndrome Study Network (NEPTUNE). NEPTUNE is part of the National Institutes of Health Rare Disease Clinical Research Network (NIH RDCRN), and is a multicenter prospective cohort study that, at the time of this study, had enrolled patients with proteinuric glomerular disease and performed comprehensive clinical and molecular phenotyping at 23 sites. Patients with secondary glomerular disease (such as diabetic kidney disease, lupus nephritis, and amyloidosis) were excluded. The overall study design has been previously published^45^. Comparable healthy renal tissue was obtained from living transplant donors (LD, n=29) and included in the study. Biopsy material from samples was micro-dissected into glomerular and tubulointerstitial compartments prior to transcriptional profiling as previously described^46^.

### Transcriptional profiling

Kidney tissue was processed as previously described^46^. Briefly, renal biopsy tissue was stored in RNAlater^®^ (ThermoFisher, Waltham, MA), and manually microdissected into tubulointerstitial and glomerular compartments. Microdissected renal biopsy specimens were processed and analyzed using Affymetrix GeneChip Human Genome U133A 2.0, U133 Plus 2.0 and Human Gene ST 2.1 Array platforms. Probe sets were annotated to Entrez Gene IDs using custom CDF version 19 generated from the university of Michigan Brain Array group^47^. For the analysis of samples profiled on the U33A 2.0 and U133 Plus 2.0 platforms, we restricted the analysis to the non-Affx probe sets common to both platforms (12,074). Expression data was quantile normalized and batch corrected using COMBAT^48^. Technical replicates of human reference RNA profiles (Stratagene, La Jolla, CA) were included with each batch and used to assess batch correction efficiency. Differential gene expression across the transcriptome was compared between patients with NS and AAV versus living donors using the SAM method^49^in MEV (version 4.9); genes were defined as differentially expressed with q-value<0.05 and fold-change thresholds were applied as defined in the manuscript. Hierarchical clustering analysis (Ward’s method) was performed in Spotfire using Z-score normalized transcript expression. CEL files and processed data used in these analyses are accessible in GEO^50^ under reference numbers: GSE104948, GSE104954, GSE108109, and GSE108112.

### Functional enrichment and network analysis

Functional networks were interrogated for common disease mechanisms and upstream regulators, shared between both diseases using: QIAGEN’s Ingenuity Pathways Analysis (IPA) (QIAGEN, Redwood City, CA, https://www.qiagenbioinformatics.com/products/ingenuitypathway-analysis). Automated assessment of literature curated cause and effect relationships was used to identify potential regulators of the shared transcriptional profile^9^. The approach derives a prediction of activation or inhibition by identifying potential network changes that can explain the observed gene expression profile (activated), or reverse the observed expression profile (inhibited). Genomatix Genome Analyzer (GGA)^51^ was used to identify pathway enrichment, and the Connectivity Map (CMap)^10,52^ and the Library of Network-Based Cellular Signatures (LINCS) Unified Environment (CLUE, http://clue.io) data version 1.0.1.1 was used to identify potential therapeutic targets through drug response signatures, and gene overexpression/knockdown signatures. At the time of analysis, LINCS contained cell line transcriptional responses to 8,878 perturbagens.

### Development of STAT1 activation gene networks and activation scores

The STAT1 activation network was generated by first developing a kidney specific STAT1 interaction network using GIANT^14^ (minimum relationship confidence of 0.70, maximal network size of 20 genes), which resulted in a 17 gene interaction network. To further refine the network, genes were assessed for functional activation in response to IFN-gamma stimulation using the Interferome database^53^, and for the presence of STAT1 and STAT family binding sites in promoter regions, resulting in a 14 gene set representing STAT1 activation. The STAT1 activation score was generated in patient samples by transforming log2 expression profiles into Z-scores, and averaging Z-scores of 14-STAT1 dependent genes to generate a STAT1 activation score for each patient sample.

### GFR estimates

GFR was estimated by the four-variable Modification of Diet in Renal Disease (MDRD) study equation^54^ for each patient in ERCB and NEPTUNE.

### Measurement of urinary EGF

uEGF concentration was measured in spot urine samples using the Human EGF Immunoassay Quantikine ELISA (R&D Systems) as previously described^17^’

### Assessment of interstitial fibrosis and tubular atrophy

Histopathology was assessed in the NEPTUNE biopsy cohort with digital images obtained from paraffin-embedded kidney tissue stained with hematoxylin and eosin, PAS, silver-based, and trichrome stains^55^. Whole slide images of glass slides from cases stored in the NEPTUNE digital pathology repository were assessed for percentage of cortex involved by IF/TA by six pathologists. The percentage of cortex involved by IF/TA was determined in each individual stain and averaged for an overall % value^56^. Cases with paired tubulointerstitial gene expression were assessed.

### Statistical analyses

Correlation analyses were performed using Pearson’s correlation within Spotfire (TIBCO). Analysis of variance (ANOVA) was used to compare PC1, PC2, and PC3 eigenvalues across FSGS, MCD, and MN in the ERCB cohort, and to assess eGFR differences across patient clusters within Spotfire (TIBCO). A two-tailed unpaired t-test (unequal variance) was used to compare eGFR differences between clusters 1 and 4.

## Author contributions

SE planned and analyzed human transcriptome data, database investigation, and correlation with clinical parameters, supervised JH, managed the project, critically reviewed data, and wrote the manuscript. VN provided critical experimental design suggestions. LM reviewed local pathology reports, and performed STAT1 outcome analysis in the NEPTUNE cohort. FE and HH performed NEPTUNE transcriptomic dataset normalization. JH contributed insight to STAT1 score development. MTL and CDC managed and provided samples from the ERCB. WJ analyzed urine EGF data. BG performed renal biopsy microdissection to obtain glomerular and tubulointerstitial compartments. HP, CSG, and JNT provided regular feedback for network defining approaches. PG provided expertise on ANCA-associated vasculitis and questions of relevance for clinicians treated patients with vasculitis. MGS provided an interpretation of eQTL results. RAL provided critical feedback on informatics approaches and expertise of JAK-STAT signaling in nephrotic syndrome. JK, PAM, and MK critically reviewed data and manuscript, and supervised the entire project. All authors contributed to and reviewed the manuscript.

## Acknowledgements

Thanks and appreciation goes to the patients who consented for their biosamples to be used in this research. The Nephrotic Syndrome Study Network Consortium (NEPTUNE), U54-DK-083912, and Vasculitis Clinical Research Consortium (VCRC), U54 AR057319, and U54 RR019497 are part of the National Institutes of Health (NIH) Rare Disease Clinical Research Network (RDCRN), supported through collaboration between the Office of Rare Diseases Research (ORDR) and the National Center for the Advancement of Translational Sciences (NCATS), part of the National Institutes of Health (NIH) Rare Disease Clinical Research Network (RDCRN), and the National Institute of Diabetes, Digestive, and Kidney Diseases. Additional funding and/or programmatic support for this project has also been provided by the Else Kröner-Fresenius Foundation (ERCB), VCRC, University of Michigan, the NephCure Kidney International and the Halpin Foundation, and the Applied Systems Biology Core at the University of Michigan George M. O’Brien Kidney Translational Core Center.

ERCB members at the time of the study: Clemens David Cohen, Holger Schmid, Michael Fischereder, Lutz Weber, Matthias Kretzler, Detlef Schlöndorff, Munich/Zurich/AnnArbor/New York; Jean Daniel. Sraer, Pierre Ronco, Paris; Maria Pia Rastaldi, Giuseppe D’Amico, Milano; Peter Doran, Hugh Brady, Dublin; Detlev Mönks, Christoph Wanner, Würzburg; Andrew Rees, Aberdeen and Vienna; Frank Strutz, Gerhard Anton Müller, Göttingen; Peter Mertens, Jürgen Floege, Aachen; Norbert Braun, Teut Risler, Tübingen; Loreto Gesualdo, Francesco Paolo Schena, Bari; Gunter Wolf, Jena; Rainer Oberbauer, Dontscho Kerjaschki, Vienna; Bernhard Banas, Bernhard Krämer, Regensburg; Moin Saleem, Bristol; Rudolf Wüthrich, Zurich; Walter Samtleben, Munich; Harm Peters, Hans-Hellmut Neumayer, Berlin; Mohamed Daha, Leiden; Katrin Ivens, Bernd Grabensee, Düsseldorf; Francisco Mampaso(†), Madrid; Jun Oh, Franz Schaefer, Martin Zeier, Hermann-Joseph Gröne, Heidelberg; Peter Gross, Dresden; Giancarlo Tonolo; Sassari; Vladimir Tesar, Prague; Harald Rupprecht, Bayreuth; Hermann Pavenstädt, Münster; Hans-Peter Marti, Bern; Peter Mertens, Magdeburg, Jens Gerth, Zwickau.

## Appendix I: Members of the Nephrotic Syndrome Study Network (NEPTUNE)

*NEPTUNE Enrolling Centers*

*Case Western Reserve University, Cleveland, OH:* J Sedor^*^ K Dell^**^, M Schachere^#^

*Children’s Hospital, Los Angeles, CA:* K Lemley^*^, L Whitted^#^

*Children’s Mercy Hospital, Kansas City, MO:* T Srivastava^*^, C Haney^#^

*Cohen Children’s Hospital, New Hyde Park, NY:* C Sethna^*^, S Gurusinghe^#^

*Columbia University, New York, NY:* G Appel^*^, M Toledo^#^

*Emory University, Atlanta, GA:* L Greenbaum^*^, C Wang^**^, B Lee^#^

*Harbor-University of California Los Angeles Medical Center:* S Adler^*^, C Nast^*‡^, J La Page^#^

*John H. Stroger Jr. Hospital of Cook County, Chicago, IL:* A Athavale^*^, M Itteera^#^

*Johns Hopkins Medicine, Baltimore, MD:* A Neu^*^, S Boynton^#^

*Mayo Clinic, Rochester, MN:* F Fervenza^*^, M Hogan^**^, J Lieske^*^, V Chernitskiy^#^

*Montefiore Medical Center, Bronx, NY:* F Kaskel^*^, N Kumar^*^, P Flynn^#^

*NIDDK Intramural, Bethesda MD:* J Kopp^*^, E Castro-Rubio^#^, E Brede^#^

*New York University Medical Center, New York, NY:* H Trachtman^*^, O Zhdanova^**^, F Modersitzki^#^, S Vento^#^

*Stanford University, Stanford, CA:* R Lafayette^*^, K Mehta^#^

*Temple University, Philadelphia, PA:* C Gadegbeku^*^ D Johnstone^**^, Z Pfeffer^#^

*University Health Network Toronto:* D Cattran^*^, M Hladunewich^**^, H Reich^**^, P Ling^#^, M Romano^#^

*University of Miami, Miami, FL:* A Fornoni^*^, L Barisoni^*^, C Bidot^#^

*University of Michigan, Ann Arbor, MI:* M Kretzler^*^, D Gipson^*^, A Williams^#^, R Pitter^#^

*University of North Carolina, Chapel Hill, NC:* P Nachman, K Gibson, S Grubbs, Anne Froment

*University of Pennsylvania, Philadelphia, PA:* L Holzman^*^, K Meyers^**^, K Kallem^#^, FJ Cerecino^#^

*University of Texas Southwestern, Dallas, TX:* K Sambandam^*^ E Brown^**^, N Johnson^#^

*University of Washington, Seattle, WA:* A Jefferson^*^, S Hingorani^**^, K Tuttle^**§^, K Klepach^#^, M Kelton^#^, A Cooper^#§^

*Wake Forest University, Winston-Salem, NC:* B Freedman^*^, JJ Lin^**^, M Spainhour^#^, S Gray^#^

*Data Analysis and Clinical Coordinating Center:* M Kretzler, L Barisoni, C Gadegbeku, B Gillespie, D Gipson, L Holzman, L Mariani, M Sampson, P Song, J Troost, J Zee, E Herreshoff, C Kincaid, C Lienczewski, T Mainieri, A Williams

*National Institute of Diabetes and Digestive and Kidney Diseases (NIDDK) Program Office:* K Abbott, C Roy

*The National Center for Advancing Translational Sciences (NCATS) Program Office:* T Urv, PJ Brooks

^*^Principal Investigator; ^**^Co-investigator^; #^Study Coordinator

^‡^Cedars-Sinai Medical Center, Los Angeles, CA

^§^Providence Medical Research Center, Spokane, WA

